# Insights Into an Antibacterial Type VI Membrane-Disrupting Effector Of Adherent Invasive *Escherichia coli*

**DOI:** 10.64898/2026.10.08.757527

**Authors:** Maïalène Chabalier, Hélène Botchorichvili, Lilly M. Billard, Eric Cascales, Nicolas Barnich, Elisabeth Billard, Laure Journet

**Affiliations:** Laboratoire d’Ingénierie des Systèmes Macromoléculaires, Aix-Marseille Université, CNRS, UMR 7255, 31 Chemin Joseph Aiguier CS7071, 13402 Marseille Cedex 09, France; M2iSH (Microbes, Intestine, Inflammation and Susceptibility of the Host), Inserm UMR 1071, Université Clermont Auvergne, INRAe USC 1382, Clermont-Ferrand, France

**Keywords:** Adherent-invasive *Escherichia coli*, T6SS, antibacterial toxin, toxin-immunity pair, membrane disruption, pore-forming, Crohn’s disease

## Abstract

Adherent Invasive *Escherichia coli* (AIEC) strains are associated with ileal Crohn’s disease and may contribute to intestinal dysbiosis through antibacterial competition. The AIEC reference strain LF82 encodes two type VI secretion systems (T6SSs), a contractile machine that injects an effector-loaded needle made of Hcp, VgrG and PAAR into neighbouring rivals, favoring niche colonization. LF82 T6SSs harbour several putative antibacterial toxin-immunity modules. Here we characterize the T6SS-1 encoded Tme^AIEC^, a PAAR-fused T6SS effector carrying a C-terminal Tme domain, hallmark of Tme (<u>T</u>ype VI <u>m</u>embrane disrupting <u>e</u>ffector) toxins, and its DUF1240-like immunity protein Tmi^AIEC^. When heterologously produced in *E. coli* K-12 as well as in *E. coli* and *Citrobacter* clinical isolates, Tme^AIEC^ is toxic only upon periplasmic targeting, and its toxicity is neutralized by Tmi^AIEC^. Periplasmically-targeted Tme^AIEC^ inserts into the inner membrane, collapses the proton motive force and increases cell permeability, consistent with a membrane-disrupting activity. Mutational analyses coupled to functional assays identified a predicted hydrophobic transmembrane segment and a conserved aspartate residue within the Tme domain as key determinants for Tme^AIEC^ toxicity. These findings establish Tme^AIEC^/Tmi^AIEC^ as a functional T6SS toxin-immunity pair and reveal conserved determinants underlying the activity of Tme-family membrane-disrupting effectors.

## INTRODUCTION

Crohn’s disease (CD) is a chronic inflammatory bowel disease affecting the ileum and colon intestinal mucosae^1^, characterized by an excessive inflammatory immune response and increased intestinal epithelial permeability. Its etiology is multifactorial and involves host genetic susceptibility, environmental factors and alteration in gut microbiota composition. CD-associated dysbiosis is characterized by reduced microbial diversity with a decrease in Bacillota (formerly called Firmicutes) and an increase relative abundance of Enterobacteriaceae. Notably, several studies have revealed that the ileal mucosa of CD patients is unusually colonized by AIEC strains compared with healthy individuals^2,3,4^. AIEC pathogenicity is characterized by their ability to adhere to and invade intestinal epithelial cells, to survive and replicate within macrophages, and to promote pro-inflammatory responses^5^. AIEC may also instigate chronic inflammation in susceptible hosts by altering the gut microbiota composition. Antibacterial competition systems are therefore of particular interest in understanding how AIEC strains colonize and persist within inflamed intestinal environments. Among these systems, the antibacterial activity of the type VI secretion system (T6SS) of various bacterial species was shown to impact microbiota composition and to promote colonization^6,7,8,9^. Interestingly, AIEC LF82 strain encodes two T6SS gene clusters^10,11^ that are expressed in digestive conditions and influence the composition of the microbiota in a mouse model of infection (unpublished results).

The T6SS is a contractile nanomachine, widespread among Gram-negative bacteria, and frequently involved in interbacterial competition. In gut pathogens and commensals, the T6SS has been shown to facilitate niche colonization by outcompeting other species. The T6SS injects effectors/toxins directly into prokaryotic target cells, and in some instances, in eukaryotic host cells. The T6SS belongs to the broad family of contractile injection systems (CISs). It is composed of a contractile tail, structurally and functionally comparable to that of contractile bacteriophages, anchored to the cell envelope by a membrane complex^12,13,14^. The tail is comprised of a baseplate^15,16,17^, which triggers polymerization of the contractile structure made of a needle surrounded by a sheath assembled in an extended conformation^18,19,20^. The needle consists of an inner tube formed by stacked Hcp hexameric rings and tipped by a spike composed of a VgrG trimer and a conical PAAR protein^21,22,23,24^. The contraction of the extended sheath propels the needle directly into the target cell. T6SS effectors/toxins can be delivered by direct association with Hcp, VgrG or PAAR, or via the help of adaptor proteins. Some effectors, called specialized effectors, are fused to Hcp, VgrG or PAAR domains for delivery^25,26,27,28,29,30^.

A large assortment of antibacterial T6SS effectors has been characterized in recent years, revealing effector families targeting essential molecules and functions in the cytoplasm (DNA, energy metabolites, protein synthesis), periplasm (peptidoglycan), or membranes of the targeted bacteria to cause cell lysis or growth inhibition^25,27,28,29,31,32,33^. These antibacterial effectors are often encoded next to cognate immunity proteins, which protect the producing cell from self-intoxication or attacks from siblings. Among periplasmic-acting effectors, two main families target the membranes: Tle (Type VI lipase effectors) phospholipases hydrolyse membrane phospholipids, whereas pore-forming or membrane-disrupting (Tme, <u>T</u>ype VI <u>m</u>embrane-disrupting <u>e</u>ffector) toxins compromise inner membrane integrity, leading to membrane depolarization, growth arrest and possibly cell death ^34,35,36,37,38,39,40,41,42^.

The recently described Tme effectors are characterized by a conserved C-terminal Tme domain that contains a predicted hydrophobic transmembrane (TM) helix flanked by a DxxK motif and a conserved aspartate residue^35^. Tme effectors are genetically associated with small inner membrane immunity proteins belonging to the DUF1240 or DUF1240-like families, termed Tmi proteins^35^. Although Tme toxins have been proposed to perturb the inner membrane of target bacteria, the molecular determinants underlying their activity remain incompletely understood.

Here we report the characterization of Tme^AIEC^, a specialized toxin with a N-terminal PAAR domain and a C-terminal Tme-domain encoded by the T6SS-1 cluster of AIEC LF82. When addressed artificially in the periplasm of *E. coli* K-12 or of clinical *E. coli* or *Citrobacter* isolates, Tme^AIEC^ production induces toxicity that it counteracted by the co-production of the Tmi^AIEC^ immunity protein. Upon periplasmic targeting, Tme^AIEC^ associates with the inner membrane, dissipates the membrane potential and increases cell permeability, consistent with membrane-disrupting activity. We further identify Asp301 and a hydrophobic TM helix of the Tme domain as key determinants of Tme ^AIEC^ toxicity. Together, these results define Tme^AIEC^/Tmi^AIEC^ as a functional T6SS toxin-immunity pair and provide mechanistic insight into the activity of Tme-family membrane-disrupting effectors.

## RESULTS

### AIEC LF82 T6SS-1 encodes a PAAR-Tme/Tmi toxin-immunity pair

The AIEC reference strain LF82 encodes two T6SS gene clusters, T6SS-1 and T6SS-2^10,11^. T6SS-1 was previously shown to encode a Tle3 phospholipase A_1 t_oxin (LF82_435) and its immunity protein Tli3 (LF82_434), located between genes encoding VgrG and PAAR proteins^43^ (Figure 1A). In addition to this Tle3 effector, T6SS-1 encodes a second putative toxin-immunity pair, comprising LF82_440 and LF82_441. *LF82_440* encodes a 418-amino-acid protein with a N-terminal PAAR_CT_2 domain (cd14744 of National Center for Biotechnology Information [NCBI] Conserved Domain Database) spanning residues 7-86) that was previously listed among Tme-family effectors^35^, in Supplementary Data 5). Indeed, LF82_440 contains a predicted Tme-domain in its C-terminal region, spanning residues 210-361 (Figure 1B,C) and shares 45% sequence identity with the Tme-domain of *V. parahaemolyticus* Tme2^35^. Consistent with the defined characteristics of Tme-family effectors^35^, this Tme-domain contains a DxxK motif, a predicted TM helix, located between residues 320 and 340, and the downstream conserved aspartate residue (Figure 1B). AlphaFold3 structural modelling of full-length LF82_440 predicted a N-terminal PAAR domain and an α-helical C-terminal region corresponding to the Tme-domain, separated by a mostly disordered region (residues 97-155) (Figure 1C). The overall confidence of the full-length model was however limited with a pTM score of 0.3 (Figure S1A). Modelling of the isolated Tme domain (residues 210–361) yielded a more confident prediction (pTM score of 0.63), and supported the presence of a predominantly α-helical fold containing the putative TM helix, and additional helices, hereafter referred to as H1 and H2 (Figure S1B, Figure 1C). The loop upstream helix H1 is predicted to comprise a β-domain composed of short β-strands. Taken together, these predictions suggest that LF82_440 is a PAAR-fused Tme-family effector. Due to the similarity to *V. parahaemolyticus* Tme2 and the conserved features of Tme-family effectors, LF82_0440 was named Tme^AIEC^.

**Figure 1.**
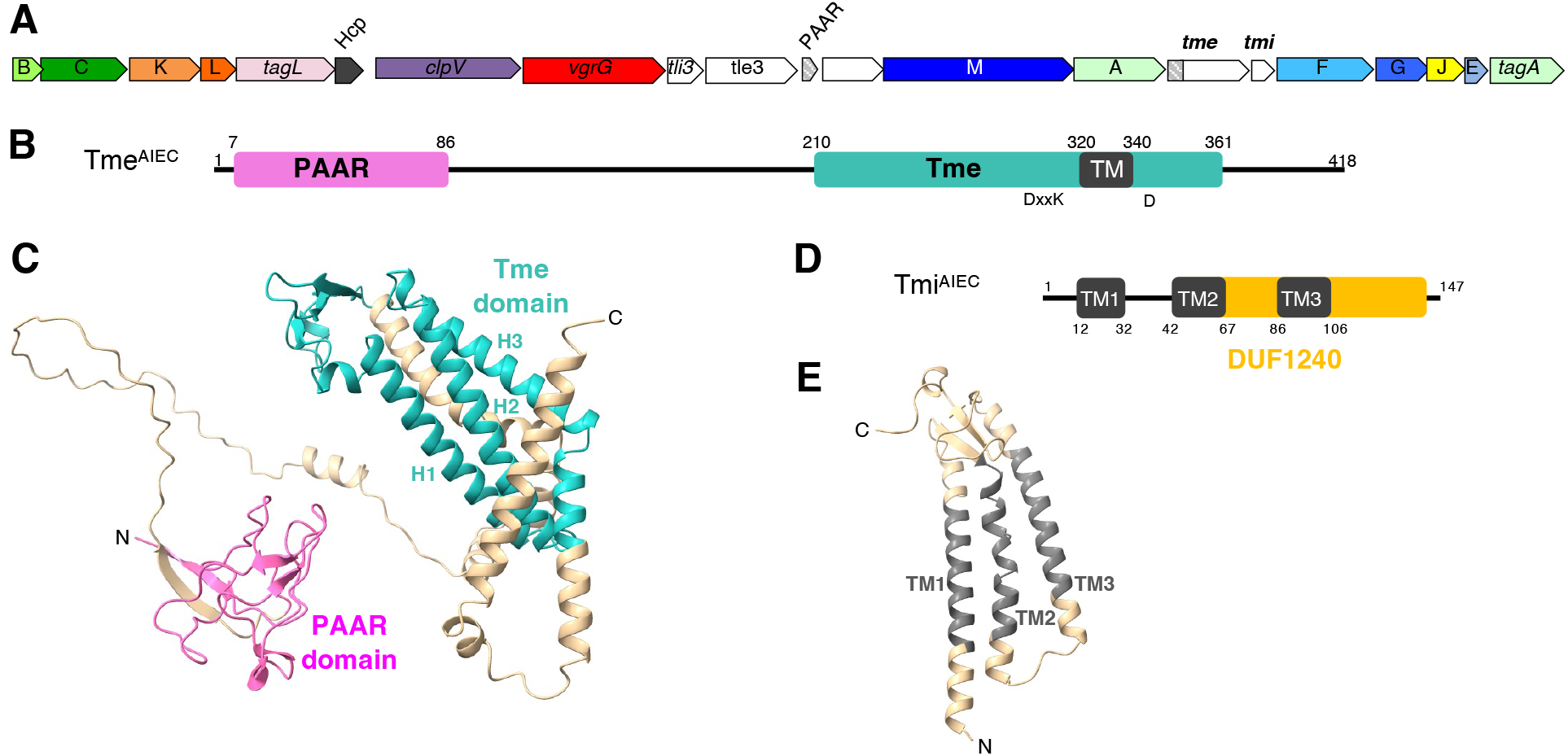
In silico analysis of LF82_440 and LF82_441, a putative Tme/Tmi couple. **A.** Schematic representation of the T6SS-1 cluster of LF82. T6SS core-components are depicted in color, identified by their Tss letter code or name, and white genes correspond to toxin/immunity modules or gene of unknown function. **B.** Schematic representation of Tme^AIEC^ (LF82_440) protein from AIEC LF82 strain. In silico PAAR detected domain (PF05488.13, PAAR_CT_2) is depicted as a pink rectangle. The Tme conserved domain (green rectangle) and conserved motif (DxxK and D, corresponding to D_301_, K_304_ and D_347_ in Tme^AIEC^) as identified in^35^ are depicted. Black rectangles correspond to TM helices predicted using TOPCONS software^69^. Numbers indicate the amino-acid number of the corresponding proteins and domains. **C.** Alphafold3 structural model of Tme**^AIEC^** (pTM=0.35, see also Figure S1A). Regions corresponding to the PAAR and Tme-domains are highlighted in pink and green, respectively. The confidence values of the model (pLDDT and PAE (Predicted Aligned Error) plot are presented in Figure S1. **D.** Schematic representation of Tmi^AIEC^ (LF82_441) protein from AIEC LF82 strain. Tmi^AIEC^, annotated as a DUF1240-containing protein, is predicted by TOPCONS software to be an integral membrane with three TM domains (amino-acids 12-32, 47-67 and 86-106). **E.** Alphafold3 structural model of the Tmi^AIEC^ (LF82_441) protein (pTM=0.79). The three TM helices are colored in grey. The confidence values of the model (pLDDT values and PAE (Predicted Aligned Error) plot are presented in Figure S1).

The downstream gene, LF82_441, encodes a small predicted inner membrane protein containing three transmembrane segments, TM1-3 (Figure 1D), forming a bundle (Figure 1E, Figure S1C). LF82_0441 is related to DUF1240-like domain proteins^35^, commonly associated with Tme-family effectors, and have been proposed to function as cognate immunity proteins. LF82_441 belonging to the DUF1240-like family, and being genetically associated to Tme^AIEC^, will thereafter be referred to Tmi^AIEC^.

### Tme^AIEC^ is toxic upon periplasmic targeting and is neutralized by Tmi^AIEC^

To test whether Tme^AIEC^ is toxic to bacteria, we heterologously produced Tme^AIEC^ in the cytoplasm or in the periplasm of *E. coli* K-12 (Figure 2AB, Figure S2). The Tme^AIEC^ coding sequence was cloned under the control of the arabinose-inducible P*araBAD* promoter of the pBAD18 vector, fused to the TorA TAT signal sequence (_TAT_-Tme) to promote periplasmic targeting, or not (Tme) for cytoplasmic production. The Tmi^AIEC^ coding sequence was cloned under the control of the IPTG-inducible P*lac* promoter of the pOK12 vector. Cytoplasmic production of Tme^AIEC^ had little effect on *E. coli* growth in spot dilution assays (Figure 2A) despite detectable production of Tme^AIEC^ by immunoblotting (Figure 2B). In contrast, periplasmic targeting of _TAT_-Tme^AIEC^ strongly impaired *E. coli* growth on agar plates, resulting in a ∼ 4-log reduction in viability compared to cells carrying with empty vectors. In liquid medium, cells producing _TAT_-Tme^AIEC^ showed a growth defect. Under these conditions, the decrease in growth was not accompanied by a drop in cell turbidity or a major loss of colony-forming units (Figure S2), suggesting that Tme^AIEC^ primarily inhibits growth (bacteriostatic) rather than causing cell lysis (bacteriolytic), as already observed for other T6SS pore-forming toxins^35,36,38,44^.

**Figure 2.**
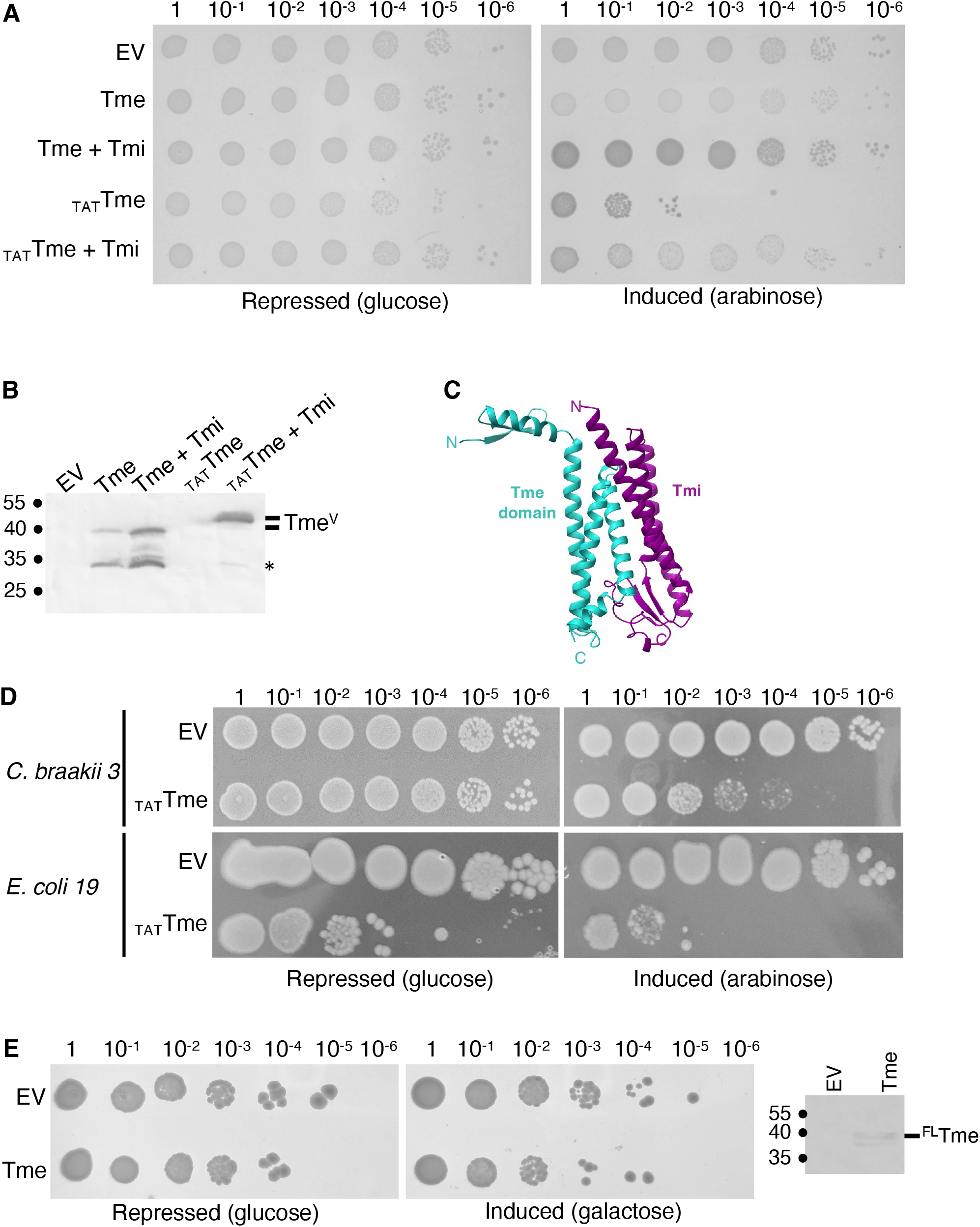
Tme^AIEC^ is toxic when produced in the periplasm of *E.* coli and Tmi^AIEC^ counteracts this toxicity. **A.** Tme^AIEC^ periplasmic toxicity is counteracted by Tmi^AIEC^. *E. coli* W3110 cells carrying empty vectors (EV, pBAD18 and pMS600) or co-transformed with pBAD18 derivative vectors allowing the production of Tme^AIEC^ (Tme, pBAD-Tme^VSVG^) or Tme^AIEC^ fused to the N-terminal TorA TAT signal peptide (_TAT_Tme, pBAD-_TAT_Tme^VSVG^) both alone (pMS600) or with the immunity protein (from pMS-Tmi^FLAG^) were serially diluted and spotted on LB agar plate containing 0.4% D-glucose or 0.2% L-arabinose to repress or induce respectively gene expression. **B.** Spots from plates (from A, induced condition) resuspended in phosphate buffer were loaded onto SDS-PAGE gel followed by an immunoblot using anti-VSVG-tag monoclonal antibodies. The molecular weight markers (in kDa) are indicated on the left. The position of VSVG-tagged Tme^AIEC^ and TAT-Tme^AIEC^ (Tme^V^ and TAT-Tme^V^) is indicated on the right. Note a slight difference of apparent migration of these two versions of Tme^AIEC^. The asterisk indicates degradation products. **C.** Alphafold3 model of the predicted complex between Tme-domain (210-361) of Tme^AIEC^ and Tmi^AIEC^ (ipTM=0.87, pTM=0.77) colored by molecules (Tme domain in turquoise, Tmi in purple). The confidence values of the model (pLDDT values and PAE plot are presented in Figure S1. **D.** Tme^AIEC^ is toxic in microbiota clinical isolates. *Citrobacter braakii* or non AIEC *E. coli* strains isolated from patients’ microbiota were used for this experiment. Cells carrying an empty vector (EV, pBAD18-TAT) or expressing Tme^AIEC^ fused to the N-terminal TorA signal peptide were spotted on LB agar plate containing 0.4% D-glucose or 0.2% L-arabinose to repress or induce gene expression, respectively. **E.** Tme^AIEC^ is not toxic when produced in S*. cerevisiae.* By4741 cells carrying an empty vector (pRS416_Gal1) or expressing Tme^AIEC^ (from pRS-^FLAG^Tme) were spotted on SDdropout (Glucose) or SGdropout (Galactose) media to repress or induce respectively gene expression. The right panel shows the production of FLAG-tagged Tme from cells spots taken from the induced plate analyzed by Western blot using anti-FLAG antibodies.

The toxicity of periplasmic Tme^AIEC^ was efficiently counteracted by the co-production of Tmi^AIEC^, as the growth cells producing both _TAT_-Tme^AIEC^ and Tmi^AIEC^ was comparable to the empty vectors control in both plate and liquid assays (Figure 2A,B, Figure S2). Taken together, these results show that Tme^AIEC^ is a periplasmic-acting antibacterial toxin and that Tmi^AIEC^ provides cognate immunity. Direct biochemical analysis of the Tme^AIEC^-Tmi^AIEC^ interaction was hindered by the aberrant migration of Tmi^AIEC^ during SDS-PAGE, which remained in the stacking gel. AlphaFold3 structural co-modeling of the Tme^AIEC^ Tme-domain with Tmi^AIEC^ predicted a stable complex with high interface confidence score (ipTM = 0.87), suggesting that the two proteins interact (Figure 2C, Figure S1D) consistent with the functional protection conferred by Tmi^AIEC^ *in vivo*. Interestingly, the presence of Tmi^AIEC^ in the AlphaFold3 co-modelling increased the confidence of the predicted Tme^−^domain structure compared with Tme^AIEC^ modeled alone (Figure S1, compare panels B and D), suggesting that Tmi^AIEC^ binding may contribute to the structural stabilization of the Tme domain.

AIEC LF82 was initially isolated from an ileal biopsy of a Crohn disease patient^45^. We therefore asked whether the Tme^AIEC^ antibacterial activity extends to other bacterial strains isolated from patient microbiota. For this, the plasmid producing _TAT_-Tme^AIEC^ was introduced into non-AIEC *E. coli* and *Citrobacter braakii* clinical isolates. Periplasmic production Tme^AIEC^ impaired the growth of all tested isolates, although the magnitude of toxicity varied between strains (2-log reduction for *C. braakii*, 2-5-log reduction for the different *E. coli* strains) (Figure 2D, Figure S3). These results indicate that Tme^AIEC^ can exert antibacterial activity in diverse Enterobacteriaceae backgrounds when delivered to the periplasm. Thus, although native T6SS-dependent delivery remains to be demonstrated, these data suggest that bacteria present in patient-associated microbiota may be susceptible to Tme^AIEC^ delivery.

Because some T6SS membrane-disrupting effectors can act on both bacterial and eukaryotic membranes, we then tested whether Tme^AIEC^ production affected the growth of the yeast *Saccharomyces cerevisiae*. Under the conditions tested, Tme^AIEC^ production did not impair yeast growth (Figure 2E) suggesting that Tme^AIEC^ does not display broad toxicity toward this eukaryotic model.

### Tme^AIEC^ becomes membrane associated upon periplasmic targeting

To get insights into the mode of action of Tme^AIEC^, we next analysed the subcellular localization of the toxin in *E. coli* W3110 (Figure 3, Figure S4). Cells producing either cytoplasmic Tme^AIEC^ or Tat-targeted Tme^AIEC^ were fractionated, and the distribution of the toxin was monitored by immunoblotting. Cytoplasmic Tme was mainly detected in the soluble cytoplasmic fraction but exhibited N-terminal degradation (Figure S4). A truncated version lacking the N-terminal PAAR domain remained soluble and led to an apparently more stable form of Tme^AIEC^ (Figure S4). In contrast, when Tme^AIEC^ was fused to the TAT signal sequence, the protein was recovered predominantly in the membrane fraction (Figure 3). To further characterize this membrane association, membrane fractions were subjected to differential extraction using high salt, urea or sodium N-lauroyl sarcosinate (SLS). _TAT_-Tme^AIEC^ was not efficiently released by high salt or urea demonstrating that it is not peripherally associated with the membranes but likely inserted in it. In addition, _TAT_-Tme^AIEC^ was solubilized by SLS, a detergent that selectively solubilizes inner membrane proteins. This behaviour indicates that periplasmically targeted Tme^AIEC^ is tightly associated with, and likely inserted into, the inner membrane (Figure 3). This inner membrane insertion from the periplasmic side upon fusion to a signal sequence, is reminiscent of the mechanism previously described for colicins^46^. Taken together, these results show that Tme^AIEC^ is soluble when produced in the cytoplasm but inserts into the inner membrane-associated after periplasmic targeting. This compartment-dependent membrane association is thus consistent with the activity of a periplasmic-acting membrane-disrupting toxin.

**Figure 3.**
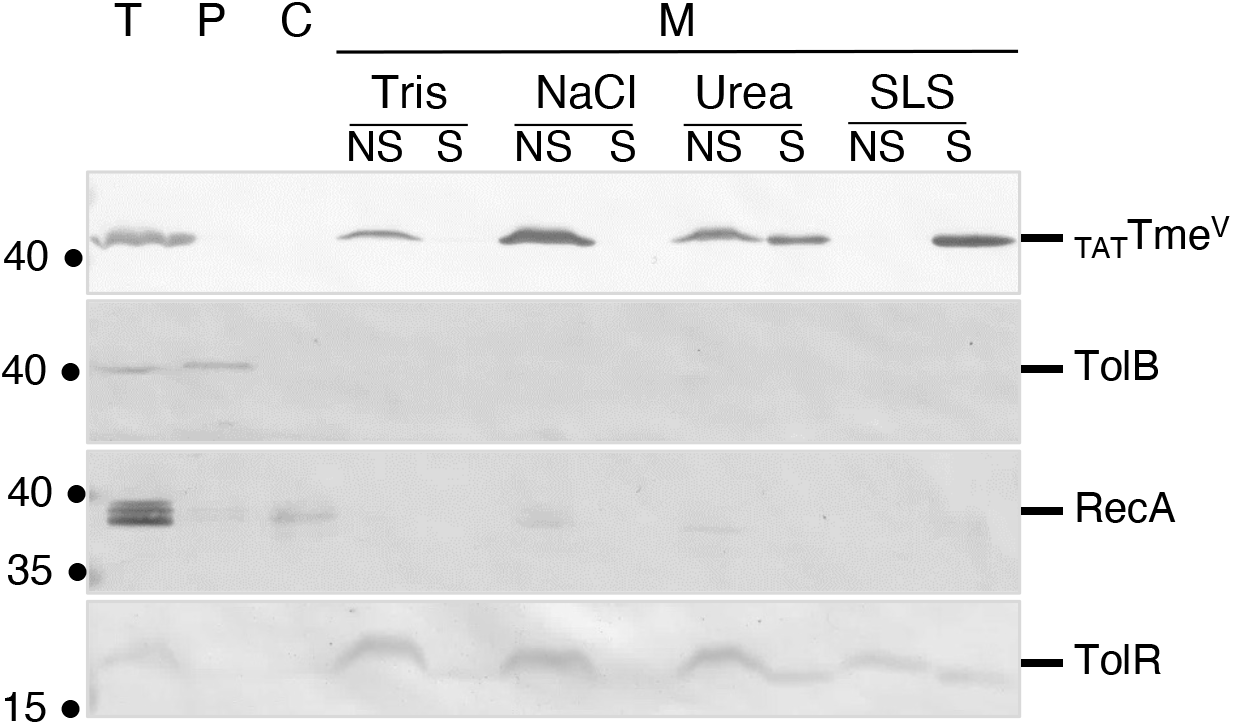
Subcellular localization of Tme^AIEC^ upon targeting to the periplasm of *E. coli*. W3110 cells carrying the pBAD18 vector expressing VSV-G-tagged Tme^AIEC^ fused with the N-terminal TorA signal peptide (TAT-Tme^V^) were fractionated to isolate the periplasmic (P), cytoplasmic (C) and membrane fractions (M). Peripheral proteins were extracted by high-salt (NaCl, PME), chaotropic agent (urea, PMH) and inner membrane-associated proteins were solubilized by N-lauroyl sarkosinate (SLS, MI). The different fractions were loaded on SDS-PAGE gels followed by an immunoblot using anti-VSV-G (TAT-Tme^V^), anti-TolB (periplasmic marker), anti-RecA (cytoplasmic marker) and anti-TolR (inner membrane marker) antibodies. The molecular weight markers (in kDa) are indicated on the left and the position of the different proteins on the right. NS, non-soluble; S, soluble.

### The predicted TM helix of Tme^AIEC^ is required for toxicity and membrane insertion

We next investigated the contribution of the predicted hydrophobic TM helix of Tme^AIEC^ to toxicity and membrane insertion. This segment, located between residues 320 and 340 (LAT**G_323_**FVA**G_327_**PLV**G_331_**VVVV**G_336_**I**G_338_**AA) is highly hydrophobic and contains several glycine residues arranged in a GxxxG pattern (GxxxGxxxGxxxxGxG, Figure 4A). These motifs, known as ‘Glycine-zippers’ are typically involved in TM helices homo-oligomerization and packing in membrane proteins and have been implicated in the assembly of transmembrane channels or oligomeric membrane complexes^47^. The helix projection of the Tme^AIEC^ TM helix and the AlphaFold3 prediction suggest that G323, G327, G331 and G338 are located on the same face of the helix (Figure 4A, Figure S5A). To test whether these glycine residues contribute to Tme^AIEC^ activity, we individually replaced Gly323, Gly327, Gly331 and Gly338 with valine. All variants were produced as TAT-fused proteins and their toxicity tested in *E. coli*. Each single substitution retained toxicity comparable to that of wild-type _TAT_-Tme^AIEC^ (Figure 4B, Figure S5B), suggesting that individual glycine residues within this motif are not essential for Tme^AIEC^ toxicity. We then assessed the role of the predicted TM segment as a whole. Deletion of amino acids 320-340 (Tme_ΔTM_) abolished _TAT_-Tme^AIEC^ toxicity (Figure 4C, Figure S5C). To determine whether toxicity depends specifically on the hydrophobic character of this segment, we generated a variant in which several hydrophobic residues were replaced by polar residues to decrease its overall hydrophobicity while preserving its helical and transmembrane features (Figure S5D,E). This variant, referred to as Tme_VVVVI_, which bears the V330N, V332Q, V333T, V334N and I337Q substitutions, was also non-toxic in *E. coli* (Figure 4C). Thus, the predicted transmembrane segment and its hydrophobic character are required for Tme^AIEC^ toxicity.

**Figure 4.**
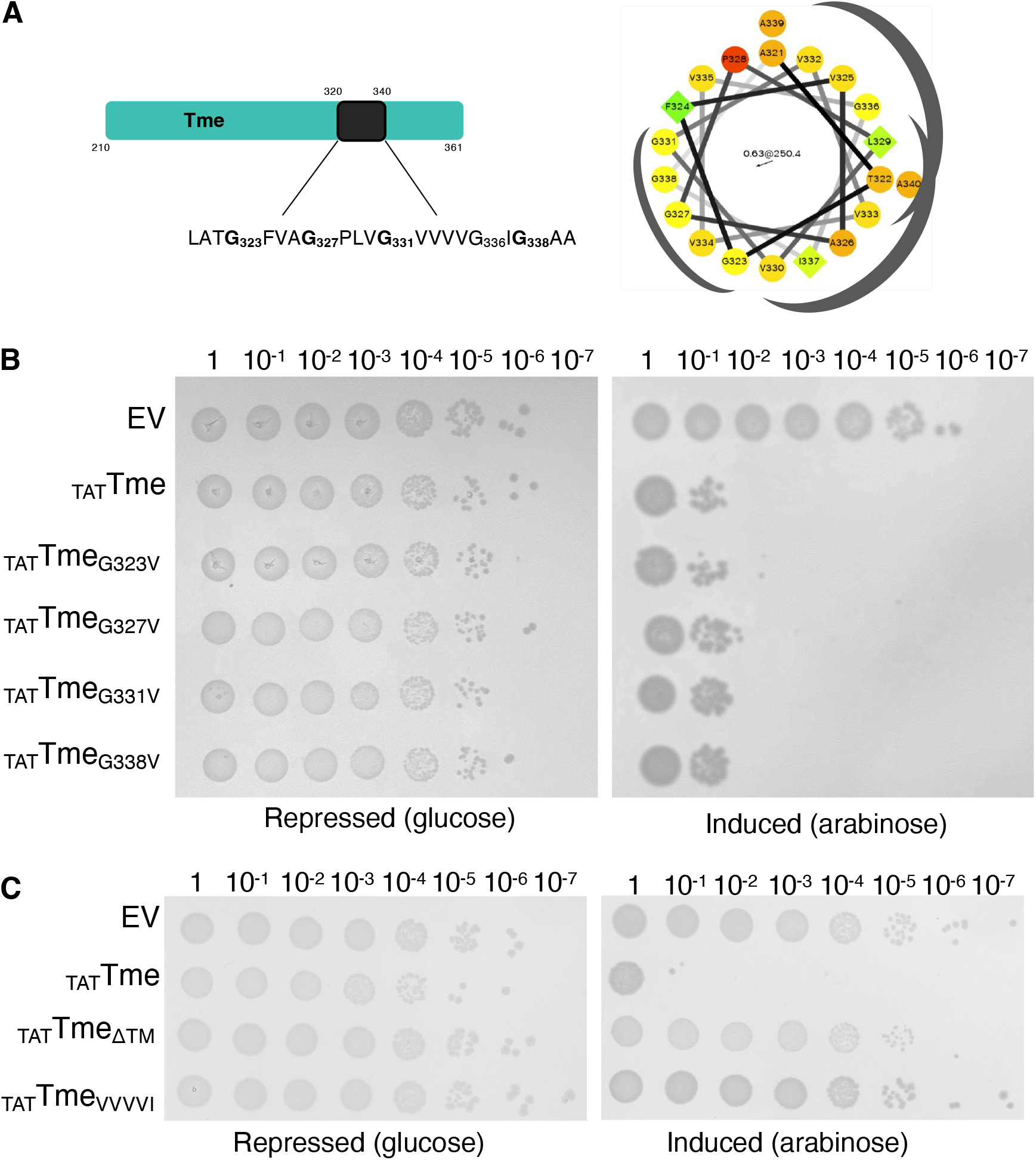
The TM helix of Tme^AIEC^ is essential for Tme^AIEC^ toxicity. **A.** Schematic representation of the Tme-domain of Tme^AIEC^ (left panel) with a putative Glycine zipper-like domain in the predicted TM helix and helix projection (right panel) highlighting the Glycine zipper-like domain and hydrophobic faces. **B.** Serial dilutions of normalized cultures of *E. coli* W3110 cells carrying pBAD18 empty vector (EV) or expressing WT or single Glycine to Valine substituted Tme^AIEC^ fused with the N-terminal TorA signal peptide (_TAT_Tme, _TAT_Tme_G323V_, _TAT_Tme_G327V_, _TAT_Tme_G331V_, or _TAT_Tme_G338V_) were spotted on LB agar plate containing 0.4% D-glucose or 0.2% L-arabinose to repress or induce gene expression, respectively. **C.** Serial dilutions of normalized cultures of *E. coli* W3110 cells carrying pBAD18 empty vector (EV), expressing periplasmic WT Tme^AIEC^ (_TAT_Tme) or Tme^AIEC^ variants: deletion of amino acids 320-340 coding sequence (_TAT_Tme_ΔTM_); simultaneous substitution of hydrophobic residues (V330N, V332Q, V333T, V334N and I337Q, _TAT_Tme_VVVVI_).

### The Asp301residue is essential for Tme^AIEC^ toxicity

In addition to the hydrophobic TM segment, Tme-family effectors, including Tme^AIEC^, contain conserved sequence features within their C-terminal Tme domain: a DxxK motif and an aspartate residue located upstream and downstream the TM, respectively^35^ (Figure S6). However, the importance of these residues has not been tested experimentally yet. To test their functional importance, we generated alanine substitutions of Asp301 and Lys204, corresponding to the DxxK motif, and of Asp347, corresponding to the conserved downstream aspartate. The D301A, K304A and D347A variants were produced as TAT-fused proteins and their toxicity was assessed in *E. coli*. Whereas wild-type Tme^AIEC^ impaired bacterial growth, the D301 variant was no longer toxic (Figure 5). In contrast, the K304A variant retained toxicity comparable to wild-type Tme^AIEC^. The D347A variant also remained toxic, although residual growth was observed at intermediate dilutions. Immunoblot analysis confirmed that the Tme^AIEC^ D301A variant was produced (Figure 5B,C), indicating that its loss of toxicity was not due to a production defect. It is noteworthy that if all constructs are detected by immunoblot when expressed in liquid medium, _TAT_-Tme wild-type and toxic mutants are not detected when cells are taken directly from induced toxicity plates (see also Figure 2B for _TAT_-Tme). Moreover, the loss of toxicity of the _TAT_-Tme^AIEC^ D301A variant was not due to impaired membrane targeting as this mutant remained associated with the inner membrane (Figure S4C).

**Figure 5.**
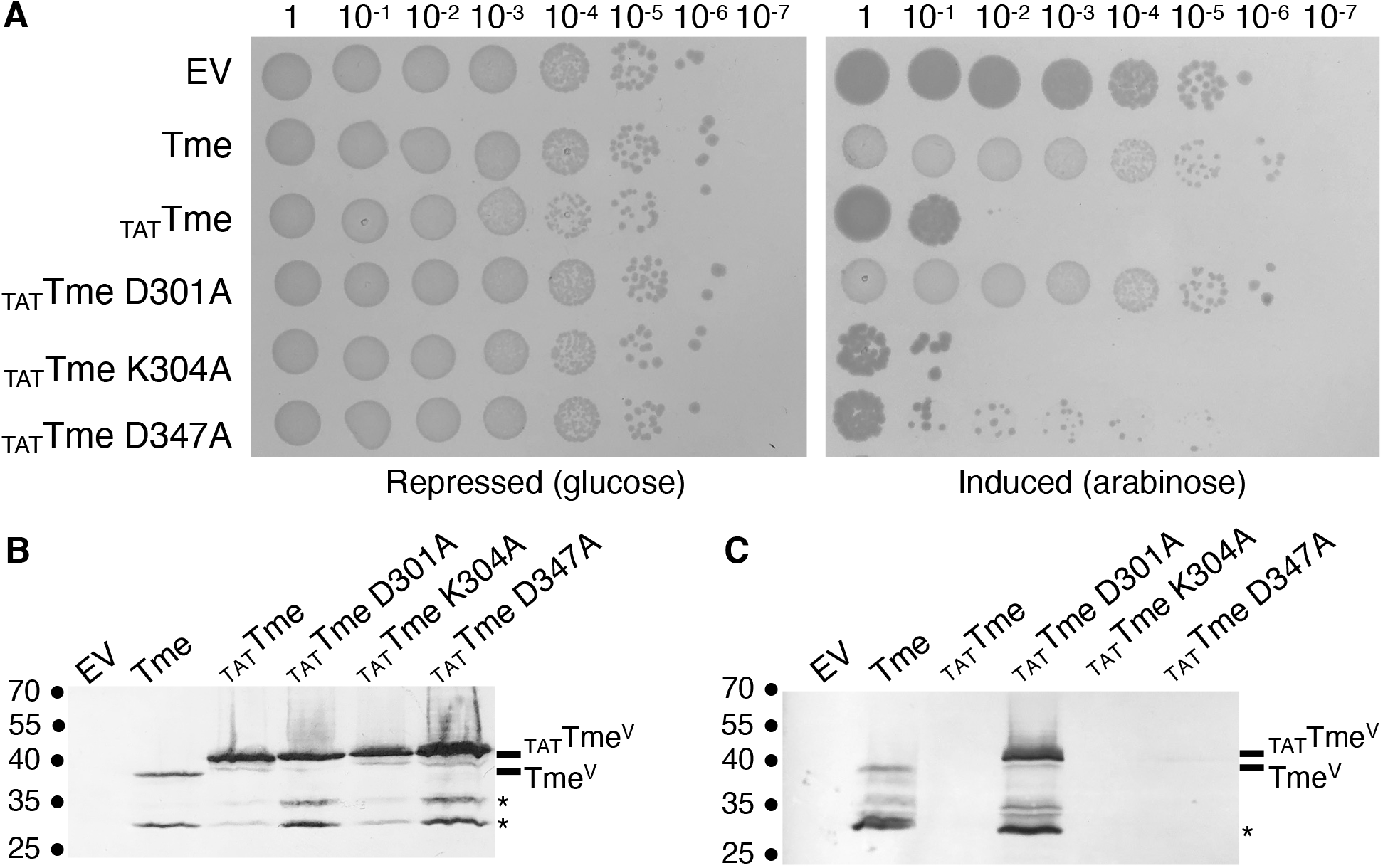
Aspartate 301 is a critical residue for Tme^AIEC^ toxicity. **A.** Serial dilutions of normalized cultures of *E. coli* W3110 cells carrying pBAD18 empty vector (EV) or expressing Tme^AIEC^ (Tme) or Tme^AIEC^ fused with TorA TAT signal peptide (_TAT_Tme), either wild type or mutated in aspartic acid 301 (D301A), lysine 304 (K304A), or aspartic acid 327 (D327A) were spotted on LB agar plate containing 0.4% D-glucose or 0.2% L-arabinose to repress or induce gene expression, respectively. **B.** Production of Tme, _TAT_Tme and _TAT_Tme mutants induced in exponential phase growth-liquid culture with of 0.2% L-arabinose were analyzed by immunoblot using anti-VSVG monoclonal antibodies. **C.** Resuspended spots from plate (from A, induced condition) were loaded onto SDS-PAGE gel followed by an immunoblot using anti-VSVG monoclonal antibodies. The molecular weight markers (in kDa) are indicated on the left. The asterisk indicates degradation products. The position of _TAT_Tme and Tme is indicated on the right. Note a slight difference of apparent migration of these two versions of Tme^AIEC^, as also observed in Figure 2B.

### Tme^AIEC^ causes permeabilization and depolarization of target cells

To test whether Tme^AIEC^ insertion in the inner membrane affects membrane integrity, we first monitored membrane depolarization using the voltage-sensitive fluorescent dye DiBAC4(3), whose fluorescence increases upon depolarization. Periplasmic targeting of wild-type Tme^AIEC^, led to an increase of the DiBAC4(3) fluorescence at 535 nm compared to control cells (Figure 6A), indicating membrane depolarization. In contrast, cells producing periplasmic-targeted Tme^AIEC^-D301A showed fluorescence levels comparable to the empty-vector control. To quantify the effect of Tme^AIEC^ on membrane integrity, we measured the transmembrane potential, Δψ, using the tetraphenylphosphonium bromide (TPP^+^) probe, whose accumulation in the cytoplasm depends on the proton motive force. Consistent with the DiBAC4(3) assay, periplasmic production of _TAT_-Tme^AIEC^ caused a marked decrease in Δψ, whereas the D301A variant had no detectable effect (Figure 6B). In both experiments, co-production of Tmi^AIEC^ with periplasmic Tme^AIEC^ restored wild-type membrane potential levels (Figures 6A and 6B). These results show that periplasmically-targeted Tme^AIEC^ dissipates the membrane potential, that Asp301 is required for this activity, and that Tmi ^AIEC^ inhibits this activity. We then tested whether Tme^AIEC^ also affects the membrane permeability using an ONPG uptake assay. ONPG is a small, 301-Da, β-galactosidase substrate that cannot penetrate intact cells. Cells producing _TAT_-Tme^AIEC^ showed increased ONPG hydrolysis, comparable to ethanol-permeabilized cells (Figure 6C). This effect was not observed upon production of cytoplasmic Tme^AIEC^, _TAT_-Tme^AIEC^ D301A, or _TAT_-Tme^AIEC^ in the presence of Tmi^AIEC^. These results indicate that Tme^AIEC^ insertion into the inner membrane increases cell permeability, altering membrane integrity and disrupting the membrane potential.

**Figure 6.**
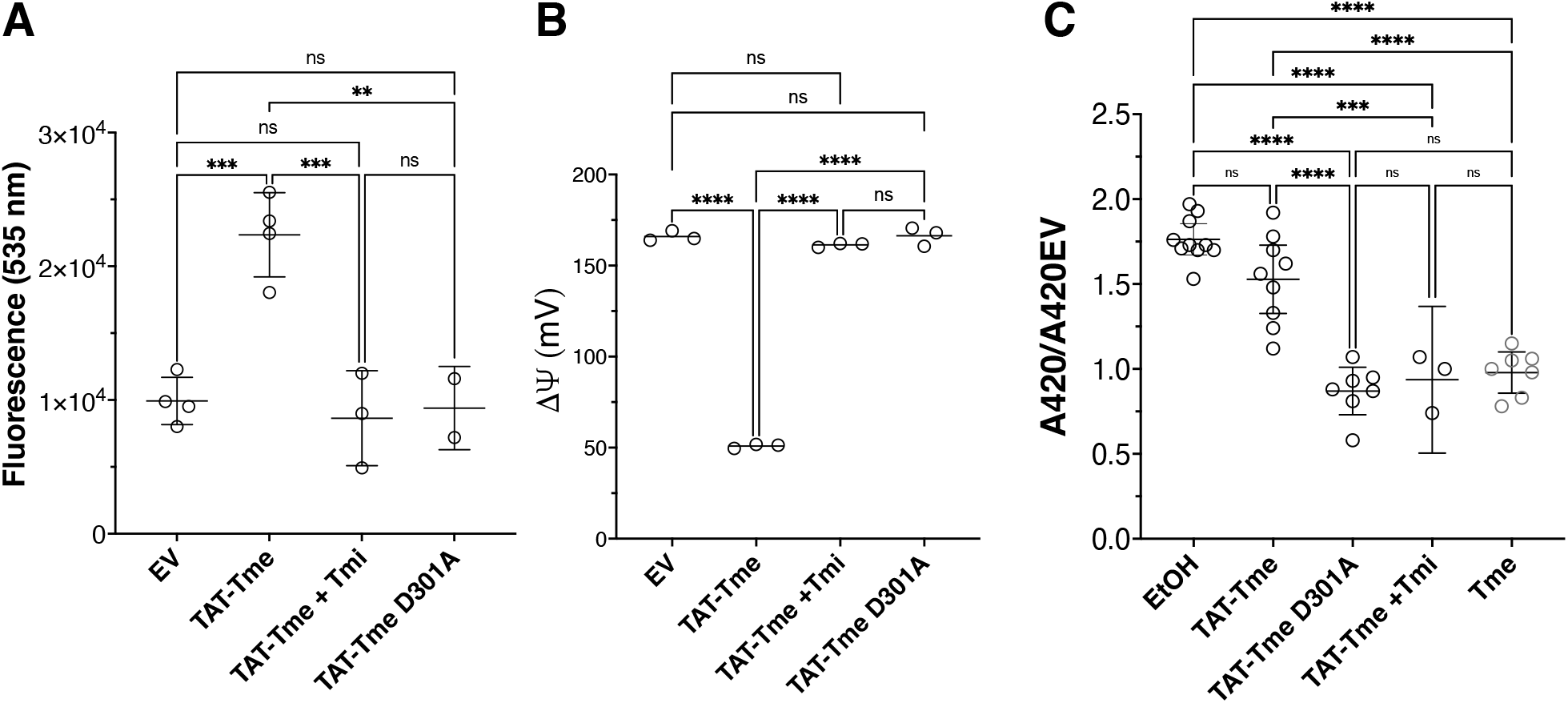
Tme^AIEC^ causes permeabilization and depolarization of target cells. **A.** Membrane potential assays were performed in *E. coli* W3110 carrying a vector (pBAD18) expressing Tme^AIEC^ or Tme^AIEC^ fused with the N-terminal TorA signal peptide (TAT-Tme) alone or with the immunity (Tmi) and a mutated version of TAT-Tme (TAT-Tme D301A). Cells were stained with DiBAC_4_(3) and fluorescence emission was measured by microplate reader (TECAN Spark®). Depolarized cells show an increase in fluorescence emitted at 535nm. Each value represents the mean of a technical triplicate. The means of two to four biological replicates are indicated. Bars show means and standard deviation. Statistical significance was calculated using ordinary one-way analysis of variance (ANOVA) followed by Tukey’s multiple comparisons test using GraphPad Prism. *** p=0.0008 (EV vs. TAT-Tme); *** p=0.0007 (TAT-Tme vs. TAT-Tme + Tmi); ** p= 0.0025; ns, p=0.9331 (EV vs. TAT-Tme + Tmi); ns, p=0.9963 (EV vs. TAT-Tme D301A); ns, p=0.9910 (TAT-Tme + Tmi vs. TAT-Tme D301A). **B.** Membrane potential of *E. coli* W3110 carrying a vector (pBAD18) expressing Tme^AIEC^ or Tme^AIEC^ fused with the N-terminal TorA signal peptide (TAT-Tme) alone or with the immunity (Tmi^AIEC^) and a mutated version of TAT-Tme^AIEC^ (TAT-Tme D301A). The electrochemical potential difference was measured by the accumulation ratio of tetraphenylphosphonium bromide (TPP+) using the Nernst equation after cell binding correction. Bars show means. Statistical significance was calculated using ordinary one-way analysis of variance (ANOVA) followed by Tukey’s multiple comparisons test using GraphPad Prism. **** p<0.0001; ns, p=0.3126 (EV vs. TAT-Tme +Tmi); ns; p=0.9981 (EV vs. TAT-Tme D301A); ns, p=0.2517 (TAT-Tme +Tmi vs. TAT-Tme D301A). **C.** Membrane permeability was tested in *E. coli* W3110 carrying the pBAD18 vector expressing Tme^AIEC^ (Tme), Tme^AIEC^ fused with the N-terminal TorA signal peptide (TAT-Tme) alone or with the immunity (Tmi) and a mutated version of TAT-Tme^AIEC^ (TAT-Tme D301A) alone. The same experiment was performed with W3110 carrying the pBAD18 empty vector. After induction with arabinose, pelleted cells were incubated with ONPG compound (301 Da) that cannot enter cells unless membrane integrity is impaired. Ethanol-treated cells were used as positive control. ONPG uptake was followed by measuring the β-galactosidase-mediated release of o-nitrophenol (A420 nm) after cell lysis. Results are expressed as ß-gal activity (A420 nm) relative to that of cells carrying the empty vectors. Bars show means and SD of at least 3 biologically independent replicates. Each value represents the mean of three technical replicates. Statistical significance was calculated using ordinary one-way analysis of variance (ANOVA) followed by Tukey’s multiple comparisons test using GraphPad Prism (GraphPad Software). **** p<0.0001; ***p=0.0002; ns, p=0.0549 (EtOH vs. TAT-Tme); ns, p=0.9826 (TAT-Tme +Tmi vs. TAT-Tme D301A); ns, p=0.9970 (TAT-Tme +Tmi vs. Tme); ns, p=0.7885 (TAT-Tme D301A vs. Tme).

## DISCUSSION

In this study, we report the characterization of the T6SS-1-encoded Tme^AIEC^/Tmi^AIEC^ toxin-immunity pair from the AIEC reference strain LF82. Tme^AIEC^ is a PAAR-fused effector containing a C-terminal Tme domain, whereas Tmi^AIEC^ is a small predicted inner membrane protein of the DUF1240-like family, encoded immediately downstream of the toxin gene. Using heterologous production in *E. coli*, we show that Tme^AIEC^ inhibits the growth of *E. coli* K-12 W3110, as well as other *E. coli* and *C. braakii* strains clinical isolates when targeted to the periplasm, and that this toxicity is neutralized by Tmi^AIEC^. Tme^AIEC^ is however not toxic when produced in yeast cells, suggesting that its activity may depend on specific features of the bacterial membrane. We further demonstrate that Tme^AIEC^ inserts into the inner membrane of *E. coli*, leading to membrane depolarization, dissipation of the ΔΨ component of the proton motive force, and increased of the cell permeability. These results suggest that Tme^AIEC^ forms large unspecific pore, in line with what was observed for *Vibrio parahaemolyticus* Tme1 and Tme2 and *Vibrio cholerae* VasX and TpeV T6SS toxins^35,44,48^. In contrast, other T6SS membrane-targeting toxins such as *S. marcescens* Ssp6/Ssp4, *P. aeruginosa* Tse4/Tse5 and *Pseudomonas putida* Tke5 have been shown to form cation selective pores^38,36,41,49^. We further identify Asp301 and the predicted hydrophobic transmembrane segment of the Tme domain as key determinants of Tme^AIEC^ toxicity. A limitation of our study is that Tme^AIEC^ activity was assessed using artificial periplasmic targeting rather than native T6SS-dependent delivery, a strategy useful to characterize antibacterial effectors whose natural delivery is difficult to reproduce under laboratory conditions. Indeed, the lack of expression of AIEC T6SS genes under laboratory conditions currently precludes *in vivo* competition assays. Therefore, while our data support the conclusion that Tme^AIEC^ intoxicates susceptible cells once delivered to the periplasm, it remains to be tested whether this effector provides AIEC a competitive advantage and contributes to niche colonization *in vivo*.

Tme^AIEC^ belongs to the class of specialized T6SS effectors because its N-terminal PAAR domain is genetically fused to the toxic C-terminal Tme domain. PAAR-domain proteins are small monomeric proteins that extends the tip of VgrG protein needle^22,50^ and are positioned at the distal end of the T6SS puncturing device, making PAAR-fused toxins ideally placed for delivery into target cells during sheath contraction. As no putative adaptor is encoded within the T6SS-1 gene cluster, this architecture suggests that the PAAR domain of Tme^AIEC^ may primarily function as a delivery module. In addition, the recruitment of Tme^AIEC^ to the T6SS needle through its PAAR domain suggests that only a single toxin is loaded and injected into the target cell per T6SS firing event. This raises the possibility that a single Tme^AIEC^ molecule is sufficient to cause membrane depolarization, without requiring toxin multimerization or multiple rounds of T6SS firing. In contrast, other small T6SS pore-forming toxins such as Tse4 and Ssp4 and Ssp6^51^, have been proposed to be loaded within the lumen of the Hcp tube, potentially allowing multiple toxin molecules to be delivered during a single firing event. This mode of delivery could, in principle, facilitate toxin oligomerization within the target cell, although oligomerization of T6SS pore-forming toxins remains poorly documented. Molecular dynamics simulations based on an AlphaFold2 structural model of Ssp4 suggested that a single Ssp4 monomer could form an ion-conducting pore^40^. The H3 TM helix of Tme^AIEC^ also contains a putative Gly-zipper motif, a feature commonly implicated in the oligomerization of transmembrane proteins. Glycine residues within such motifs can promote helix-helix packing, and a previous study showed that substituting the glycine residues of the Gly-zipper motif in the last TM helix of the T6SS pore-forming toxin Tse4 reduced antibacterial activity^36^. In contrast, mutation of the corresponding Gly-zipper residues in Tme^AIEC^ did not impair toxicity, suggesting that these residues are not essential for Tme^AIEC^ activity. These glycines are not conserved among Tme^AIEC^ homologs, including *V. parahaemolyticus* Tme2 (Figure S6), further arguing against an essential role for this motif in Tme-family toxin oligomerization or pore formation. Consistent with the hypothesis that Tme^AIEC^ acts as a monomer, AlphaFold3 multimer predictions for full-length Tme^AIEC^ or the isolated Tme-domain yielded models with very low confidence scores (ipTM < 0.12) for dimer, trimer, tetramer, or hexamers. Biochemical characterization of Tme^AIEC^ will be thus required to determine whether the protein can oligomerize and to establish whether oligomerization contributes to its membrane-disrupting activity.

Pore-forming toxins initially fold into a water-soluble, typically monomeric structure. Following recruitment to the target membrane, soluble pore-forming toxins undergo large structural rearrangements and/or oligomerization that promote membrane insertion and pore formation^52,53^. For example, once translocated in the periplasm, the C-terminal domain of pore-forming colicins such as colicin A, which comprises 10 α-helices in its soluble state, undergoes a major structural rearrangement. This process starts with the insertion into the membrane of a hydrophobic hairpin formed by helices H8–H10, which is buried in the soluble structure, followed by rearrangement of the remaining helices to form a transmembrane ion channel^54,55,56, 57,58^. Our data support a model in which Tme^AIEC^ inserts into the inner membrane after reaching the periplasmic compartment as a soluble protein. Tme^AIEC^ is produced in the cytoplasm of the attacker cell in a soluble, non-toxic, conformation. Upon delivery in the periplasm of the target cell, we propose that Tme^AIEC^ undergoes a structural transition allowing its insertion in the inner membrane via at least TM helix H3. In addition to the TM helix identified by most TM predictors (H3, aa 320-340), MEMSAT-SVM, a supervised learning method that predicts the likelihood of a TM helix participating in pore formation, also identified two additional helices present in the AlphaFold3 model of Tme-domain of Tme^AIEC^ (H1, aa 265-281; and H2, aa 295-315) (Figure 1C, Figure S1B, Figure S7), suggesting that the Tme-domain may comprise a larger transmembrane domain (TMD). Whether the Tme TMD or putative pore-forming domain is restricted of H3 only or also includes H1 and H2 remains to be determined. Our results clearly demonstrate that H1 and H2 alone are insufficient to drive membrane insertion. Conversely, reducing the hydrophobicity of H3 impaired Tme^AIEC^ toxicity, suggesting that H3 hydrophobicity may play a specific role in pore formation. Our model is not only consistent with the mechanism of pore-forming colicins, but also with the recent cryo-EM structure of a soluble form of the Ptx2 T6SS pore-forming toxin in which the putative hydrophobic TMD— thought to mediate membrane insertion—is buried inside the protein^39^. Interestingly, the AlphaFold3 model of Ptx2 suggest that the helices of this TMD are exposed, suggesting that the AlphaFold model represents the membrane-inserted form of the toxin^39,40^. Our observations are however insufficient to establish a mechanism for the formation of the pore by Tme^AIEC^. Structural characterization of soluble and membrane-inserted Tme^AIEC^ will therefore be important to define the architecture of the Tme domain and provide further insights into its membrane-disrupting mechanism. The role of Asp301, that we have shown to be required for Tme^AIEC^ toxicity, needs also to be understood. Asp301 is part of the DxxK motif conserved in the H2 helix of Tme domains (Figure S7). Asp301 is not required for insertion in the inner membrane and probably contributes to a later step such as formation or stabilization of a membrane-permeabilizing state. In addition, the partial phenotype observed for the D347A variant suggests that the downstream conserved aspartate may contribute to optimal activity, but is not strictly essential for Tme^AIEC^ toxicity.

Finally, we have shown that Tme^AIEC^ toxicity is neutralized by the Tmi^AIEC^ immunity protein (Figure 2). Tmi^AIEC^, which belongs to the DUF1240-like family, is predicted to be an integral 3-TM inner membrane protein with a predicted in/out/in/out topology (Figure 1E). Most of the identified Tmi proteins were indeed predicted to possess 3-6 TM helices^35^ and have been proposed to bind cognate Tme proteins to prevent their insertion, oligomerization or formation of the pore. Although we could not biochemically demonstrate a direct interaction between Tme^AIEC^ and Tmi^AIEC^, a confident AlphaFold3 model (ipTM=0.87, Figure S1D) predicts a plausible interface between Tmi^AIEC^ and the Tme domain of Tme^AIEC^ including the H1-H3 helices of Tme^AIEC^ and the 3 TM segments and C-terminal region of Tmi^AIEC^ (Figure S8). It is worthy to note that Tmi^AIEC^ therefore mediate contacts with the two determinants of Tme^AIEC^ toxicity identified in this work, H3 and the H2 Asp301 residue. Immunity proteins to pore-forming colicins are also membrane proteins that prevent pore formation^59,60,61^. One may hypothesize that Tmi^AIEC^ immunity proteins work similarly by preventing productive insertion of Tme^AIEC^ into the inner membrane, by blocking toxin oligomerization or pore formation, or by stabilizing an inactive toxin conformation. Future experiments aimed at detecting the Tme^AIEC^-Tmi^AIEC^ complex in native membranes or mapping the interaction interface are exciting avenues for future research.

Further biochemical experiments and mutagenesis will be required to determine the molecular mechanism underlying the membrane disrupting activity/pore formation of Tme proteins and T6SS pore-forming toxins in general. Nevertheless, these results expand our knowledge of this still poorly characterized pore-forming family.

## MATERIAL AND METHODS

### Bacterial strains, plasmids and growth conditions

Bacterial strains and plasmids used in this study are detailed in Supplementary Table S1. AIEC *E. coli* LF82 chromosomal DNA was used as genomic template for PCRs. *Escherichia coli* strain DH5α was used for cloning procedures. *E. coli* W3110 was used for transmembrane potential measurements, fractionation and permeability assays. *E. coli* W3110 and Saccharomyces cerevisiae By4741 were used for toxicity tests, as well as *Citrobacter brakii* and non-pathogenic *E. coli* isolates obtained from mucosa-associated microbiota of patients enrolled in clinical studies (respectively METABIOTE^62^ and CEALIVE^63^. *E. coli* strains were routinely grown in LB medium at 37°C and *S. cerevisiae* By4741 were grown in YPAD at 30°C with shaking. Plasmids were maintained by the addition of ampicillin (100 μg/mL) or kanamycin (50 μg/mL). Yeast transformants were selected on Synthetic Dextrose Minimal Medium (SD) medium without uracile. Expression of genes from pBAD vectors was induced with L-arabinose 0.2% (and repressed with Glucose 0.4 %). Expression of genes in Yeast was induced on Synthetic Galactose minimal medium (SG).

### Plasmid construction

Custom oligonucleotides were synthesized by Sigma Aldrich or IDT (Integrated DNA Technologies) and are listed in Supplementary Table S1. Polymerase chain reactions (PCR) were performed using a Biometra thermocycler using the Q5 High fidelity DNA polymerase (New England Biolabs). Plasmids pBAD_TAT_-Tme^VSV-G^, were constructed by restriction-ligation cloning. Plasmids pMS-Tmi^FLAG^, pBAD-Tme^VSV-G^, and pBAD-TmeΔPAAR^VSV-G^ were constructed by restriction-free cloning^64^ as previously described^65^. Briefly, genes of interest were amplified with oligonucleotides introducing extensions annealing to the target vector. The double-stranded product of the first PCR has then been used as oligonucleotides for a second PCR using the target vector as template. The Gly323-to-Val, Gly327-to-Val, Asp301-to-Ala, Lys304-to-Ala, Asp327-to-Ala substitutions were introduced in the pBAD-Tme and pBAD_TAT_Tme plasmids, by QuickChange PCR-based targeted mutagenesis. Mutations were confirmed by DNA sequencing (Eurofins).

### Bacterial toxicity assays

W3110 strain and clinical isolates (*E. coli*, *C. brakii*) carrying appropriate plasmids were grown in LB at 37°C for 16 h. Bacterial suspensions were normalized to an OD_600_ of 1, serially diluted in PBS and 8µL drops of each dilution were spotted onto selective LB agar plates containing 0.4% D-glucose or 0.2% L-arabinose to repress or induce gene expression, respectively.

### Saccharomyces cerevisiae toxicity assays

BY4741 Yeast cells transfected with pRS416_Gal1 and its derivative encoding Tme^AIEC^ were grown in selective SD lacking uracil at 30°C for 16h. Yeast suspensions were normalized to an OD_600_ of 1, serially diluted and 8µL drops of each dilution were spotted onto selective SD agar or SG agar plates, to repress or induce gene expression, respectively.

### Fractionation assay

Fractionation assay have been performed as previously described^65,66^. SDS-Polyacrylamide gel electrophoresis was performed using standard protocols. For immunostaining, proteins were transferred onto 0.2 mm nitrocellulose membranes (Amersham Protran), and immunoblots were probed with primary antibodies (see below) and goat anti-mouse secondary antibodies coupled to alkaline phosphatase and developed in alkaline buffer in presence of 5-bromo-4-chloro-3-indolylphosphate and nitroblue tetrazolium.

### Cytoplasmic membrane permeability assays

Membrane permeability assays using 2-nitrophenyl-β-D-glactopyranoside (ONPG, 301.3 Da) have been performed as previously described^36^ and have been adapted. Briefly, overnight cultures of bacterial strains were diluted to obtain OD_600_=0.05 in LB medium, grown to mid-log phase and induced with 0.02% (w/v) L-arabinose for 1h. Bacterial cells equivalent to 1mL at OD_600_=0.5 were collected by centrifugation and resuspended in 1mL of potassium phosphate buffer 0.05mM pH7,4 containing 2.4 mM ONPG (SIGMA life science). Cells were incubated at 37 °C for 1.5h, reaction was stop by adding sodium carbonate 0.33M. The cells were broken by adding 0.07% sodium dodecyl sulfate (5µL SDS 20%) and 0.3 % chloroforme (5 µL chloroforme 100%) and centrifuged to remove unbroken cells or fragments. ONPG uptake was followed by measuring the intracellular β-galactosidase-mediated release of o-nitrophenol (at A=420 nm).

### Measurements of transmembrane potential

#### DiBAC_4_(3) assay

Membrane permeability assay have been performed as previously described^36,38^ and have been adapted. Briefly, overnight cultures of bacterial strains were diluted to obtain OD_600_=0.05 in LB medium, grown to mid-log phase and induced with 0.02% (w/v) L-arabinose for 1h. Cells were pelleted by centrifugation and the OD_600_ set to 0.5. Cell suspension (100μL) were added to a staining mixture containing 1 µg.mL^−1^ DiBAC_4_(3) (Eurogentec) and incubated at room temperature for 20min in the dark. Samples were analyzed on a microplate reader (TECAN Spark®). Fluorescence excitation was performed at 485 nm and fluorescence emission was measured at 535nm.

#### PMF measurements

The electrochemical potential difference was measured by the accumulation ratio of tetraphenylphosphonium bromide (TPP^+^) using the Nernst equation after cell binding correction, as previously described^67,68^ with modifications. Cells were grown in LB until OD =0.8 and induced with 0.2% of L-arabinose for 1 hour. Cells were pelleted by centrifugation and resuspended into 100 μL of Tris-HCl 100 mM pH7.8, EDTA 1mM. After incubation for 3 min at 37°C, cells were diluted 30 folds into Sodium phosphate buffer (NaPi) 100 mM pH7.8, pelleted by centrifugation and resuspended into 30 μL of NaPi 100 mM pH7.8. Cells were kept on ice, then diluted 25-fold into NaPi 100 mM pH7.8 supplemented with 1 mM of KCl, 0.4% of glucose. [^3^H]-TPP^+^ (Amersham) was diluted into 0.5 mM TPP^+^ (Sigma-Aldrich) in order to have ∼ 25,000 cpm per μL. 2 μL of the TPP^+^ solution were added into 98-μL aliquots of the cell suspension. As control for non-specific binding of TPP^+^, 98-μL aliquots of the cell suspension were treated with 10 μM of carbonyl cyanide *m*-chlorophenylhydrazone (CCCP, Sigma-Aldrich) for 10 min prior to the addition of the TPP^+^ solution. After incubation for 10 min at 37°C, cells were recovered by filtration on GF/F filters (Whatman) and washed three times with 2 mL of NaPi 100 mM pH 7.8. Incorporated [^3^H]-TPP^+^ were measured using a Hidex 300 SL automatic TDCR liquid scintillation counter and the transmembrane potential was calculated as described by the Nernst equation.

### SDS-PAGE and Western blot analyzes

Equivalent of 0.6 OD_600_ units of cell culture for *E. coli* and 1 OD_600_ unit for *S. cerevisiae* was loaded on SDS-PAGE and transferred onto 0.2µm nitrocellulose membrane (Amersham protran). Anti-VSV-G (SIGMA #V5507), anti-FLAG (clone P5D4, Sigma-Aldrich), anti-TolB (periplasmic marker, laboratory collection polyclonal antibodies), anti-RecA (cytoplasmic marker, abcam ab63797), anti-EF-Tu (cytoplasmic marker, Roche), anti-IscS (cytoplasmic) or anti-TolR (inner membrane marker, laboratory collection polyclonal antibodies) were used as primary antibodies and anti-mouse phosphatase alkaline-conjugated (Interchim #115-055-003) or anti-rabbit phosphatase alkaline-conjugated (Interchim #111-055-003) were used as secondary antibodies. Revelation was performed in alkaline buffer (pH9) using 5-bromo-4-chloro-3-indolyl phosphate/nitroblue tetrazolium in the presence of 10 mM MgCl2.

## Acknowledgments

We thank Maximilien Rouzaud, Nicolas Rivard and the members of the Cascales research group for insightful discussions. We thank Moly Ba, Melissa Guilbert, Fabien Gomez and Audrey Gozzi for technical assistance. We thank Mathilde Bonnet, Anthony Buisson and Michaël Rodrigues for providing *E. coli* and *Citrobacter* clinical isolates, and the Agence Nationale de la Recherche (16-IDEX-0001 CAP 20-25) that funded METABIOTE and CEALIVE studies. This work was supported by the Aix-Marseille Université (amU), the Centre National de la Recherche Scientifique (CNRS) and grant from the Agence Nationale de la Recherche (ANR-18-CE15-0013).

## Author contributions

LJ, EB, NB and MC designed the experiments; MC, HB, LB, EC and LJ conducted the experiments, collected and analyzed the data; MC and LJ wrote the article; EB and EC critically revised and edited the manuscript.

