## Supplementary information for "Insights Into an Antibacterial Type VI Membrane-Disrupting Effector Of Adherent Invasive *Escherichia coli*"

This PDF includes:

- Figures S1 to S8

- References for Supplementary information

**Figure S1**

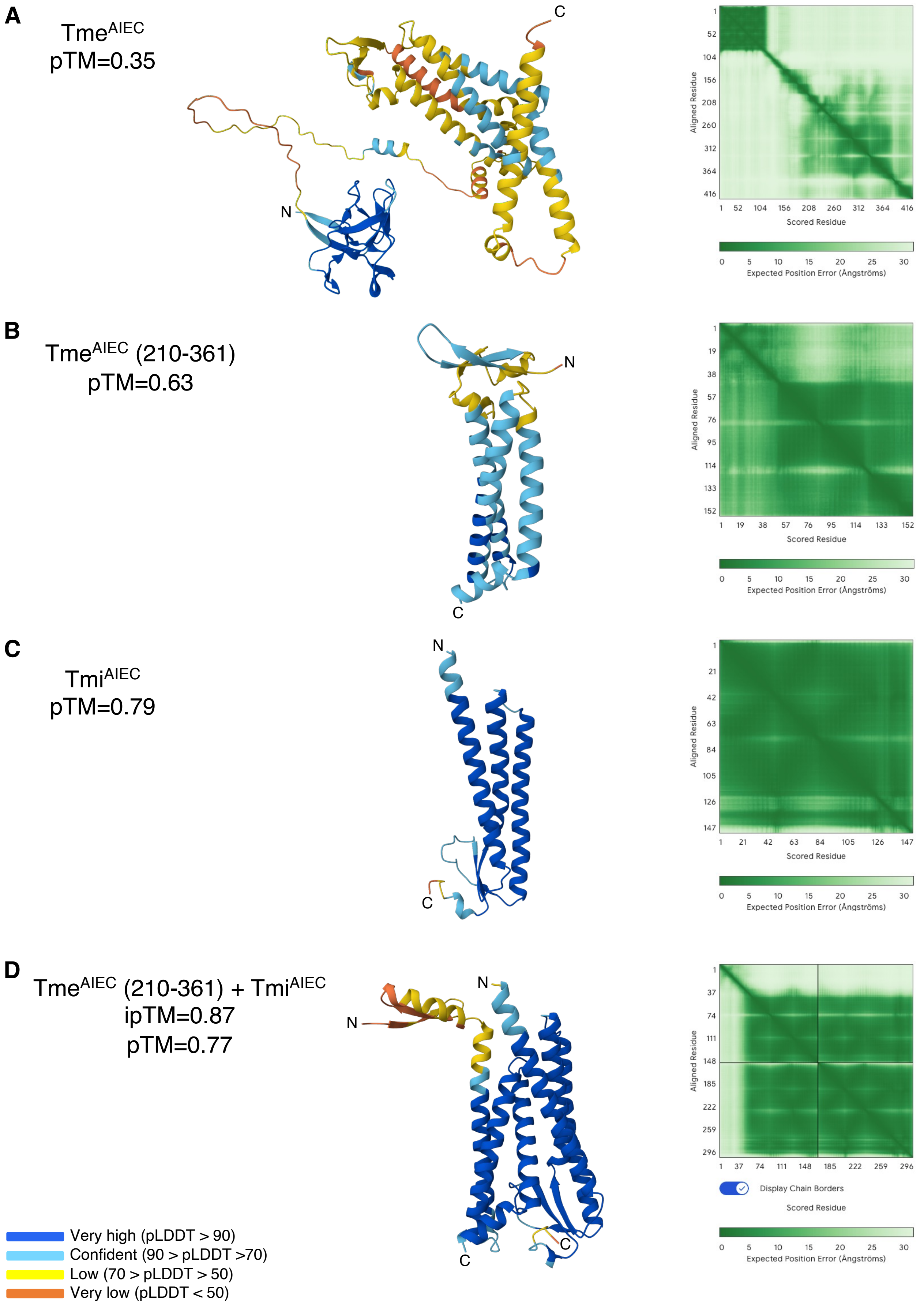
A
Tme$^{AIEC}$
pTM=0.35
C
N
1
52
104
156
208
260
312
364
416
Aligned Residue
Scored Residue
0 5 10 15 20 25 30
Expected Position Error (Ångströms)
B
Tme$^{AIEC}$ (210-361)
pTM=0.63
N
C
1
19
38
57
76
95
114
133
152
Aligned Residue
Scored Residue
Expected Position Error (Ångströms)
C
Tmi$^{AIEC}$
pTM=0.79
N
C
1
21
42
63
84
105
126
147
Aligned Residue
Scored Residue
Expected Position Error (Ångströms)
D
Tme$^{AIEC}$ (210-361) + Tmi$^{AIEC}$
ipTM=0.87
pTM=0.77
N
N
C
C
1
37
74
111
148
185
222
259
296
Aligned Residue
Display Chain Borders
Scored Residue
Expected Position Error (Ångströms)
Very high (pLDDT > 90)
Confident (90 > pLDDT >70)
Low (70 > pLDDT > 50)
Very low (pLDDT < 50)

**Figure S1. AF3-predicted structure of (A) Tme$^{AIEC}$, (B) Tme-domain of Tme$^{AIEC}$ (amino-acids 210-361), (C) Tmi$^{AIEC}$ and (D) of the Tme$^{AIEC}$ (210-361)/Tmi$^{AIEC}$ complex.**

AF3-structural models are colored by pLDDT value (left panels), and the corresponding PAE (Predicted Aligned Error) plots are presented on the right. pLDDT confidence values: dark blue, very high; blue, confident; yellow, low; orange, very low confidence.

### Figure S2

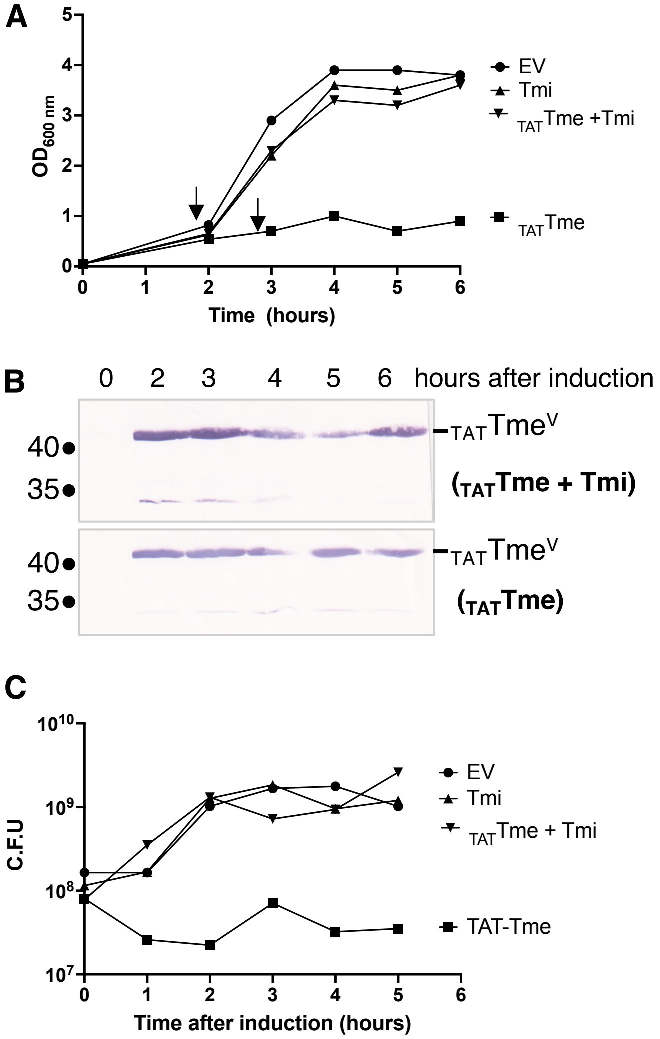


**Figure S2. $Tme^{AIEC}$ periplasmic toxicity leads to growth inhibition but not to cells lysis.**

**A**. Growth curve of *E. coli* W3110 cells carrying empty vectors (EV, pBAD18 and pMS600) or co-transformed with pBAD18 derivative vectors allowing the production of $Tme^{AIEC}$ (Tme, pBAD-$Tme^{VSVG}$) or $Tme^{AIEC}$ fused to the N-terminal TorA TAT signal peptide ($_{TAT}Tme$, pBAD-$_{TAT}Tme^{VSVG}$) both alone (pMS600) or with the immunity protein (from pMS-$Tmi^{FLAG}$). Cells were cultivated in LB in the presence of glucose 0.4 % until OD600 reach 0.5, washed and gene expression was induced with 0.2% arabinose (arrows). This experiment was repeated three times with similar results and a representative curve is shown.

**B**. Samples from the bacterial growth culture from **A** were taken at the different indicated time points after induction to analyze $_{TAT}Tme^{AIEC}$ ($_{TAT}Tme^{V}$) production by Western blot using anti-VSV-G antibodies. The molecular weight markers (in kDa) are indicated on the left and the position of $_{TAT}Tme^{V}$ is shown on the right.

**C**. Samples from bacterial growth culture from **A** were taken at the different indicated time points after induction to determine colony forming units (CFU).

### Figure S3

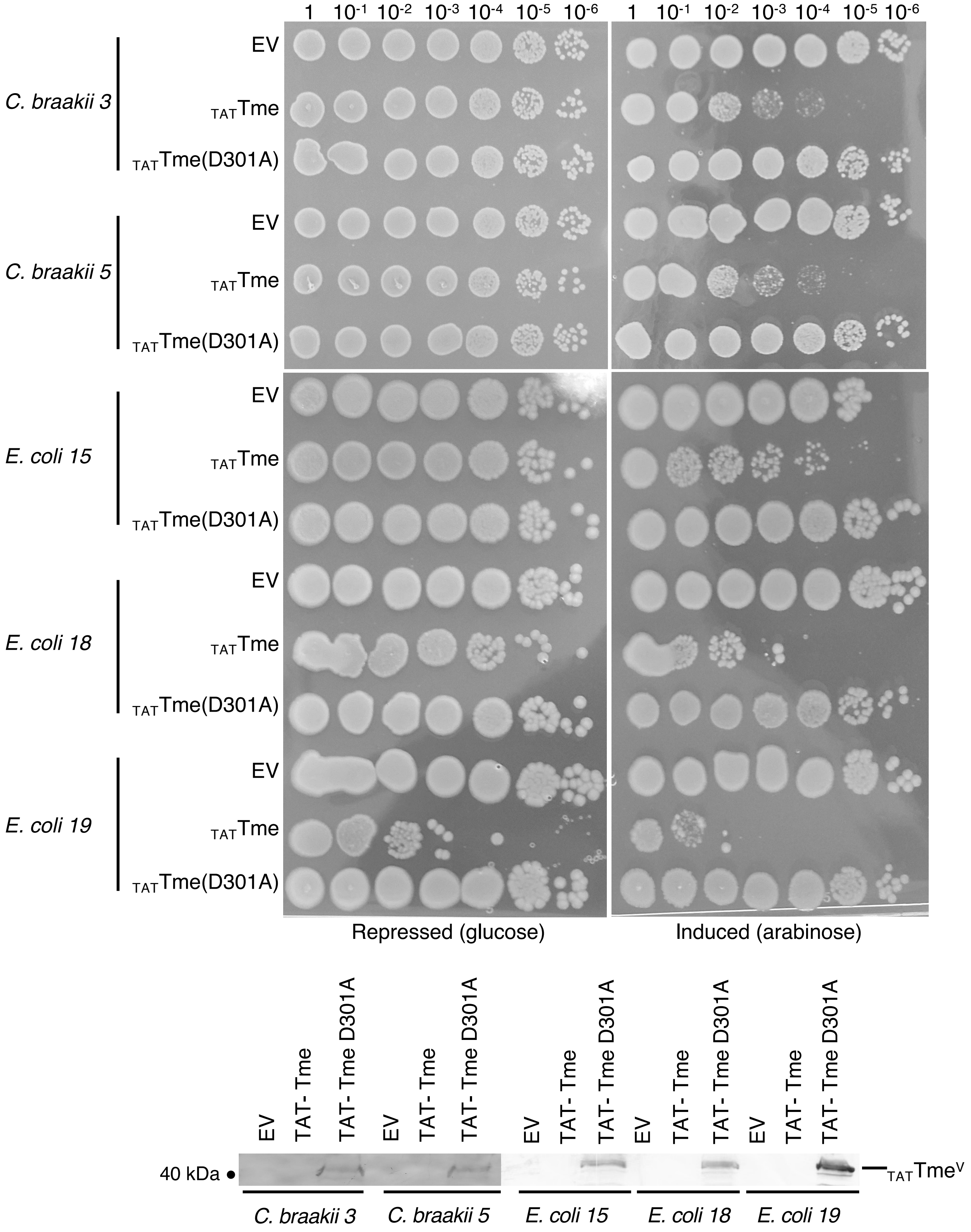
1
10-1
10-2
10-3
10-4
10-5
10-6
C. braakii 3
C. braakii 5
E. coli 15
E. coli 18
E. coli 19
EV
TATTme
TATTme(D301A)
Repressed (glucose)
Induced (arabinose)
EV
TAT- Tme
TAT- Tme D301A
40 kDa
TATTmeV

**Figure S3. TmeAIEC is toxic in microbiota clinical isolates.**

Different *Citrobacter braakii* or non AIEC *E. coli* strains isolated from patients' microbiota were used for this experiment. Cells carrying an empty vector (EV, pBAD18-TAT) or expressing $Tme^{AIEC}$ or $Tme^{AIEC}$ (D301A) mutant fused to the N-terminal TorA signal peptide were spotted on LB agar plate containing 0.4% D-glucose or 0.2% L-arabinose to repress or induce gene expression, respectively. Spot from plates, resuspended in phosphate buffer, were loaded onto SDS-PAGE gels followed by an immunoblot using anti-VSV-G ($_{TAT}$TmeV, $_{TAT}$TmeD301A (lower panel).

**Figure S4**

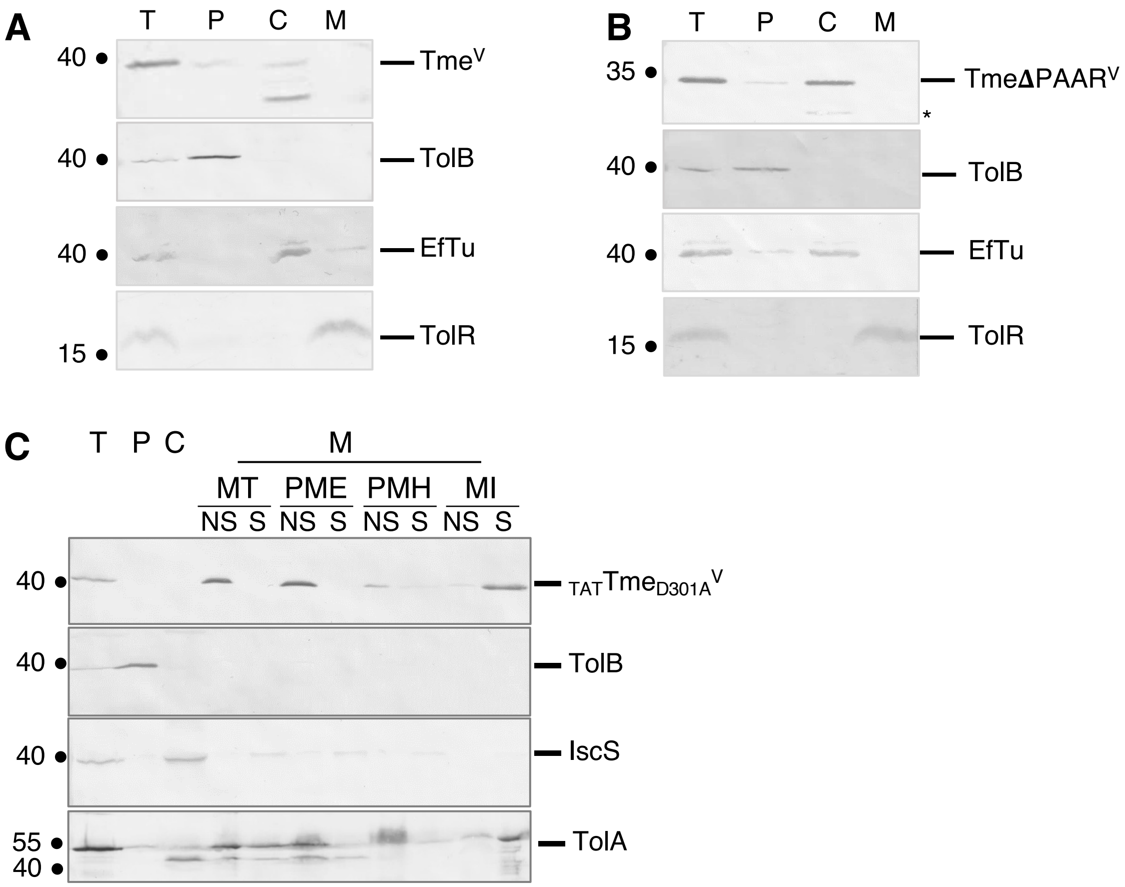


**Figure S4. Subcellular localizations of Tme$^{AIEC}$, Tme ΔPAAR and $_{TAT}$TmeD301A.**

**A**. *E. coli* W3110 cells carrying a pBAD18 vector expressing VSVG-tagged Tme$^{AIEC}$ (Tme$^{V}$) were fractionated to isolate the periplasmic (P), cytoplasmic (C) and membrane fractions (M). The different fractions were loaded onto SDS-PAGE gels followed by an immunoblot using anti-VSV-G (Tme$^{V}$), anti-TolB (periplasmic marker), anti-RecA (cytoplasmic marker), anti-EFTu (cytoplasmic marker), anti-IscS (cytoplasmic marker) and TolR (membrane marker) antibodies. The position of the different proteins and of a degradation product of Tme$^{V}$ (*) is indicated on the right. The molecular weight markers (in kDa) are indicated on the left. Note the difference of migration between Tme and $_{TAT}$Tme that may be due either to the uncleaved TAT signal sequence or a different migration profile between the two forms.

**B**. Same experiment as in **A** with *E. coli* cells carrying a pBAD18 vector expressing VSVG-tagged-Tme deleted from its first 86 amino acids (TmeΔPAAR$^{V}$).

**C**. Same experiment as in **A** with *E. coli* cells carrying a pBAD18 vector expressing VSVG-tagged-TmeVSV-G-tagged Tme$^{AIEC}$D301A mutant fused with the N-terminal TorA signal peptide ($_{TAT}$TmeD301AV). T, total cells; NS, non-soluble; S, soluble, MT, Tris; PME, Tris NaCl; PMH, urea; MI, SLS.

#### Figure S5

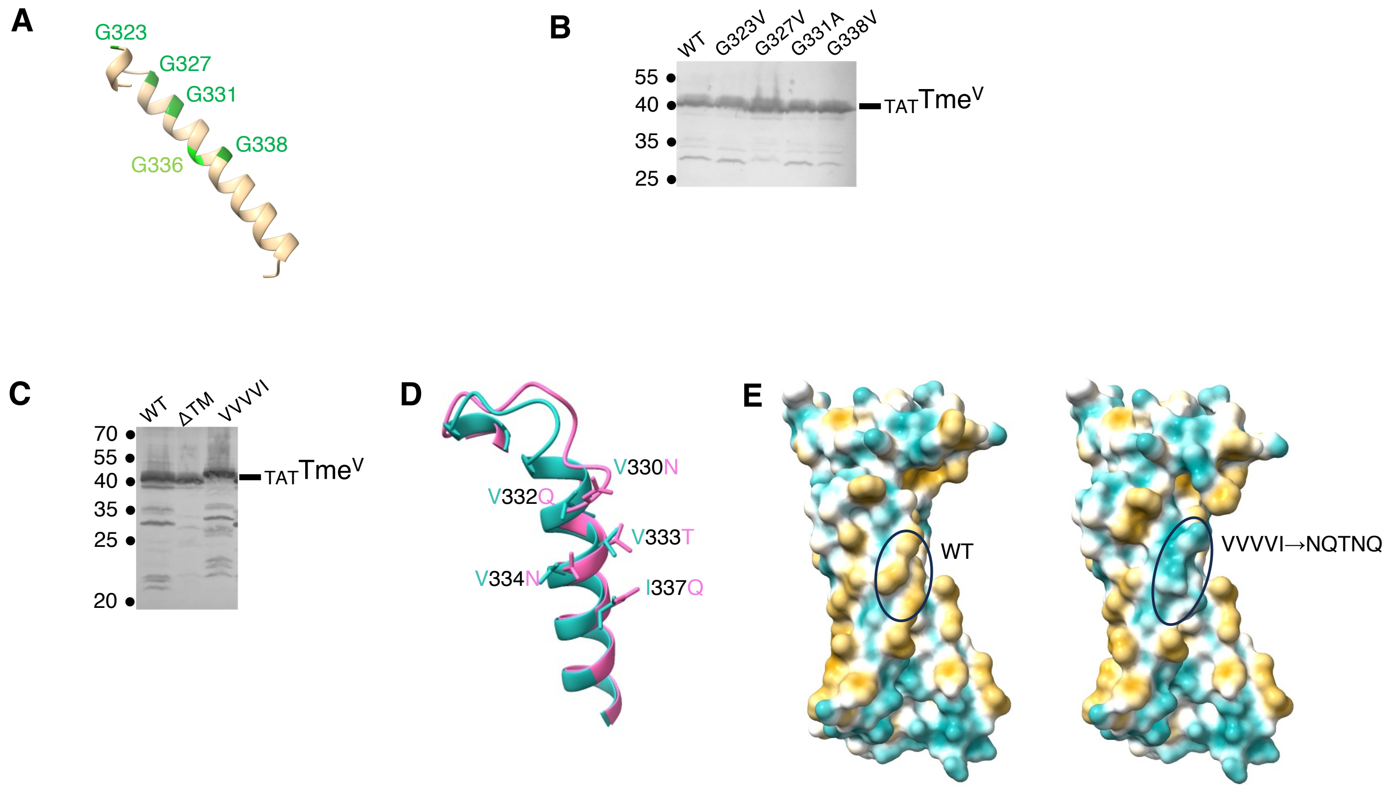


**Figure S5. The TM helix of Tme$^{AIEC}$ is an essential determinant for Tme$^{AIEC}$ toxicity.**

**A**. AF3 model of Tme-domain, showing only the region a.a 320-350, where glycines are highlighted in green. G323, G327, G331 and G338 are on the same face.

**B**. Glycine-zipper analysis. Production of $_{TAT}$Tme and $_{TAT}$Tme indicated G to V mutants induced in exponential phase growth-liquid culture with of 0.2% L-arabinose were analyzed by immunoblot using anti-VSV-G monoclonal antibodies.

**C**. TM helix analysis. Production of $_{TAT}$Tme, $_{TAT}$TmeΔTM and $_{TAT}$TmeΔVVVVI mutants induced in exponential phase growth-liquid culture with of 0.2% L-arabinose were analyzed by immunoblot using anti-VSV-G monoclonal antibodies (left panel).

**D**. AF3 model of the Tme-domain of WT Tme$^{AIEC}$ and VVVVI-NGTNQ mutant were superimposed. Mutated residues are highlighted (WT Tme-domain in green, VVVVI-NGTNQ mutant in pink). Only a.a 320-348 of the Tme-domain is shown.

**E**. AF3 models of the Tme-domain of WT Tme$^{AIEC}$ (left) and VVVVI-NGTNQ mutant (right) highlighting the modification of hydrophobicity in the mutant.

### Figure S6

**A**

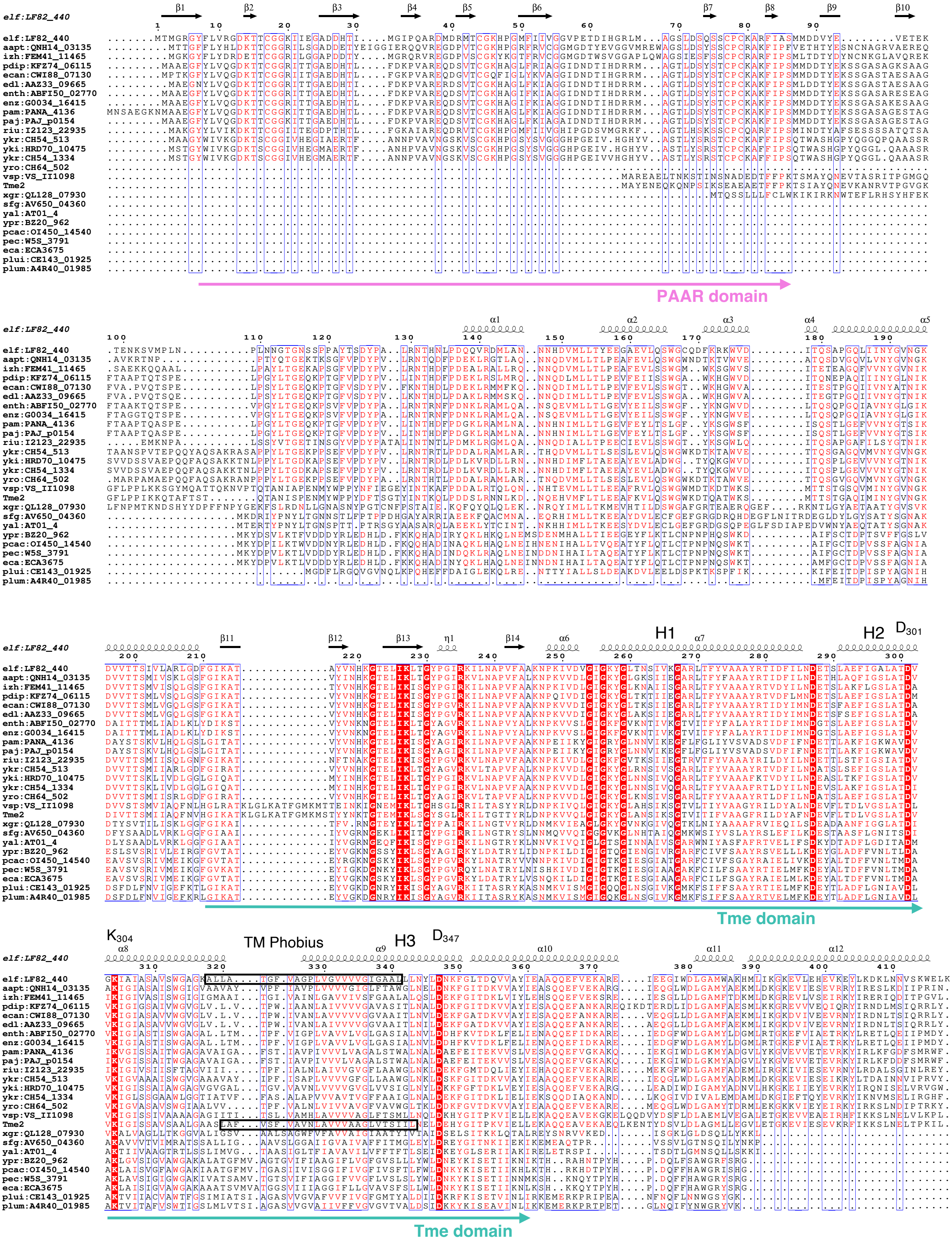
elf:LF82_440
aapt:QNH14_03135
izh:FEM41_11465
pdip:KFZ74_06115
ecan:CWI88_07130
edl:AAZ33_09665
enth:ABFI50_02770
enz:G0034_16415
pam:PANA_4136
paj:PAJ_p0154
riu:I2123_22935
ykr:CH54_513
yki:HRD70_10475
ykr:CH54_1334
yro:CH64_502
vsp:VS_II1098
Tme2
xgr:QL128_07930
sfg:AV650_04360
yal:AT01_4
ypr:BZ20_962
pcac:OI450_14540
pec:W5S_3791
eca:ECA3675
plui:CE143_01925
plum:A4R40_01985
PAAR domain
Tme domain
TM Phobius
H1
H2
H3
D301
K304
D347

**Figure S6. In silico analysis of Tme$^{AIEC}$.**
Multiple sequence alignment of Tme$^{AIEC}$ (elf:LF82_440) orthologs detected using BlastP analysis against the KEGG GENES database. Selected non redundant sequences were aligned using MultAlin (Corpet et al., 1988). The sequence of Tme2 of Vibrio parahaemolyticus (Tme2, WP_047706523.1, Fridman et al., 2020) was included in the alignment. Ordered locus names of the identified genes are indicated on the left. The conserved residues are white on a red background. The semi-conserved residues are red, and the non-conserved residues are black. The green arrow represents the Tme-domain. The position of the DxxK motif and the D347 conserved residues is indicated, as well as the putative TM detected by Phobius for Tme$^{AIEC}$ and Tme2 are indicated by a rectangle. Figure prepared using MultAlin (Corpet et al., 1988) and ESPript (Robert & Gouet, 2014).

**Figure S7**

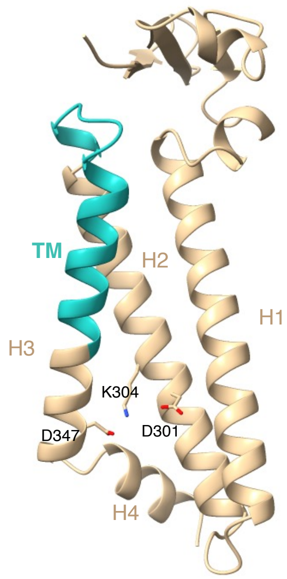


**Figure S7. AF3-structural model of the Tme-domain** (a.a 210-316) highlighting conserved residues (D301, K304 and D347) and the predicted TM helix (TM, in green).

### Figure S8

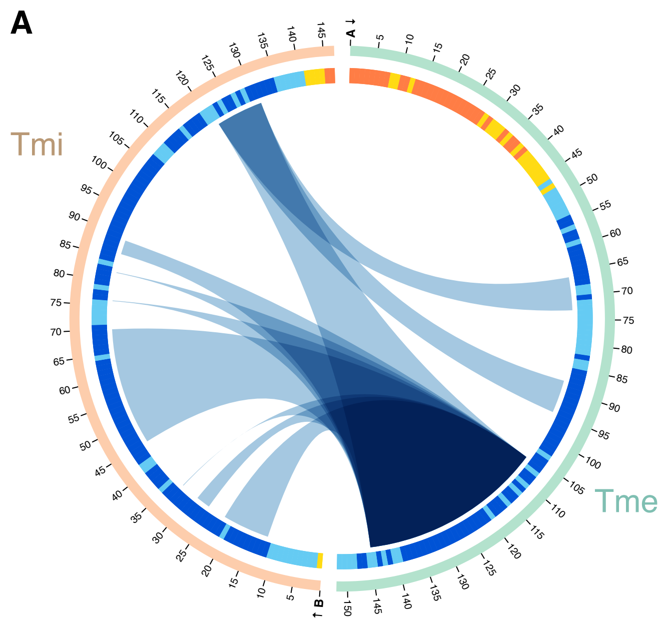


| Tme domain | | Tmi |
|---|---|---|
| Protein-Protein [1] | | piCSi: 0.92 |
| **Link** | **Sequence 1** | **Sequence 2** |
| L1 | A – 66-73 | B – 121-131 |
| L2 | A – 88-95 | B – 121-131 |
| L3 | A – 106-145 | B – 12-22 |
| L4 | A – 106-145 | B – 26-29 |
| L5 | A – 106-145 | B – 33-33 |
| L6 | A – 106-145 | B – 45-70 |
| L7 | A – 106-145 | B – 76-76 |
| L8 | A – 106-145 | B – 82-82 |
| L9 | A – 106-145 | B – 86-89 |
| L10 | A – 106-145 | B – 121-131 |

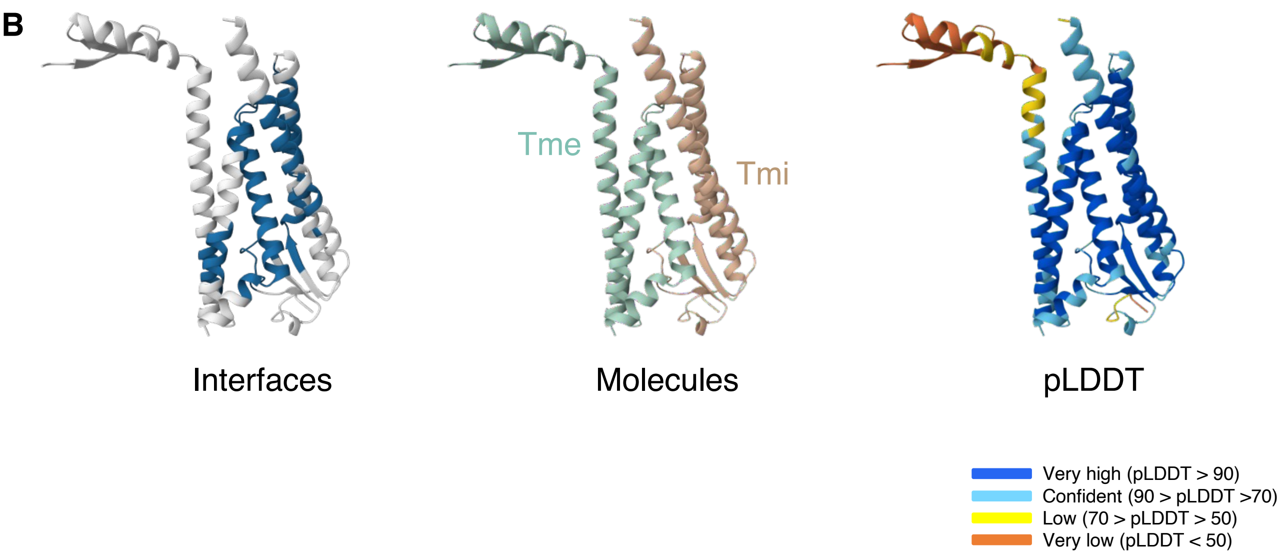


**Figure S8. Alphabridge analysis of Tme$^{AIEC}$(210-361)-Tmi$^{AIEC}$ predicted interface.**
**A**. Alphabridge representation of Tme$^{AIEC}$ (210-361)-Tmi$^{AIEC}$ complex model presented in Figure S1D (Álvarez-Salmoral et al., 2024). Confidence level for interface detection was set on Default.
**B**. AF3-structural model of the complex is are colored by interfaces (in blue, left panel), by molecules (Tme-domain in green, TmiAIEC in beige, middle panel), and by pLDDT confidence values: dark blue, very high; blue, confident; yellow, low; orange, very low confidence (right panel).
